# Discovery and characterization of llama VHH targeting the RO form of human CD45

**DOI:** 10.1101/2020.09.01.278853

**Authors:** Deepti Rokkam, Patrick J. Lupardus

## Abstract

CD45 is an abundant and highly active cell-surface protein tyrosine phosphatase (PTP) found on cells of hematopoietic origin. CD45 is of particular importance for T-cell function, playing a key role in the activation/inactivation cycle of the T-cell receptor signaling complex. The extracellular domain of CD45 is comprised of an N-terminal mucin-like domain which can be alternatively spliced to a core domain (RO) consisting of four domains with fibronectin 3 domain (FN3)-like topology. The study of CD45 has been hampered by a small set of publicly available antibodies, which we characterized as specific to the N-terminal FN3 domains of CD45 RO. To broaden the human CD45 reagent set, we identified anti-CD45 single domain VHH antibodies from a post-immune llama phage display library. Using a yeast display domain mapping system and affinity measurement we characterized seven unique clonotypes specific for CD45 RO, including binders that target each of the four FN3-like domains. These VHH molecules are important new tools for studying the role of CD45 in T-cell function *in vitro* and *in vivo*.

## Introduction

CD45/PTPRC is a highly abundant and promiscuous cell-surface protein tyrosine phosphatase (PTP) found on cells of hematopoietic origin that has a primary role in dephosphorylating immunoreceptor tyrosine-based activation/inactivation motifs (ITAM/ITIMs) in order to regulate the activity of lymphocytes (Hermiston et al., 2003). CD45 is of particular importance for T-cell function, playing a key role in both activation and inactivation of T-cell receptor (TCR) signaling (Alexander, 2000). On the activation side, CD45 activates the Lck kinase by dephosphorylating an autoinhibitory Tyr residue early in the TCR activation cycle (Sieh et al., 1993). Later, “kinetic exclusion” of CD45 molecules from the TCR synapse allows a buildup of ITAM phosphorylation and activation of TCR signaling (Carbone et al., 2017; James and Vale, 2012). Importantly, CD45 has also been shown to dephosphorylate tyrosine-based phosphorylation sites on cytokine receptors (Irie-Sasaki et al., 2001), suggesting a general role in homeostatic regulation of phosphorylation in lymphocytes.

CD45 is a uniquely structured Type I membrane protein, with a large (~600 amino acid) N-terminal extracellular domain coupled to tandem intracellular phosphatase domains. The extracellular region contains an N-terminal mucin-like domain encoded by exons 1, 2, and 3 (also called A, B, and C) which can be alternatively spliced to a core domain consisting of four domains with FN3-like topology (Chang et al., 2016). The mucin-like domain contains multiple O-linked glycosylation sites (Thomas, 1989), while the core RO form, consists solely of the four core domains, contains a number of N-linked glycosylation motifs (Hermiston et al., 2003). As a result, the size and structure of the CD45 extracellular domain is highly heterogeneous between both individual protein monomers on the surface of the cell, as well as between unique cell types expressing different CD45 splice forms.

Single domain antibodies, also known as VHH (<u>VH</u> domain on a <u>H</u>eavy chain) antibodies, are found in a number of higher eukaryotic species, including the camelid family and sharks (Muyldermans, 2013). Antibody discovery and engineering of VHH obtained from llamas, alpacas, and camels has become an important part of the antibody engineering toolkit, both for discovery of research and diagnostic tools as well as next generation antibody therapeutics (Meyer et al., 2014). VHH (when removed from the Fc domain) have the advantage of small size (~12-14 kD) and high stability, which allows access unique therapeutic spaces when compared to a standard monoclonal IgG format (Ingram et al., 2018). Combined with simple phage display discovery methods that do not require heavy/light chain pairing, straightforward humanization, and simple manufacture makes VHH and attractive platform for drug discovery, particularly when developing bispecific tools or therapeutics.

In order to advance our understanding of CD45 function, we set out to discover and characterize both mAb and VHH-based binders for the human CD45 RO splice form encompassing the D1 to D4 FN3 like domains, which are found on all CD45-expressing cells. We sequenced and expressed several human CD45 antibodies from available hybridoma cell lines, as well as executed a llama VHH discovery campaign to identify new single domain binders to human CD45. One major goal of the study was to identify the CD45 domains targeted by the antibodies to select for molecules that bound the RO form. We developed a yeast display domain mapping system for screening candidate binders and characterized their affinity by Surface Plasmon Resonance (SPR). From this work, four antibodies and seven new VHH clonotypes were identified that bind across all four domains of CD45 RO extracellular domain (ECD) with a range of affinities.

## Results/Discussion

CD45 is a commonly used cell surface marker for characterization of T cell subsets by flow cytometry. In particular the RO form has been used to identify activated and memory T cell populations (Hermiston et al., 2003). Our initial discovery efforts were focused on characterizing publicly available hybridoma cell lines that target the RO form of human CD45, in particular the D1 to D4 FN3-like domains (Fig 1A). We identified three hybridomas, gap8.3, 4B2, and 9.4, as well as a fourth antibody called VHE from the patent literature (Kolbinger, 2017). For the hybridoma lines, we sequenced the VH and VL domains from the hybridoma and cloned the sequences into the parent mouse IgG2A backbone. For the VHE antibody, we cloned the VH and VL sequences into the human IgG1 backbone. We also generated scFv molecules by pairing the VH and VL domains with an 18 amino acid Gly-Ser linker. Recombinant forms of the IgG antibodies as well as scFvs linked to a human IgG1 Fc were then generated by transient expression in expi293 cells followed by purification on protein A resin. These IgG and scFv molecules were used for domain mapping experiments and biophysical analysis.

**Figure 1:**
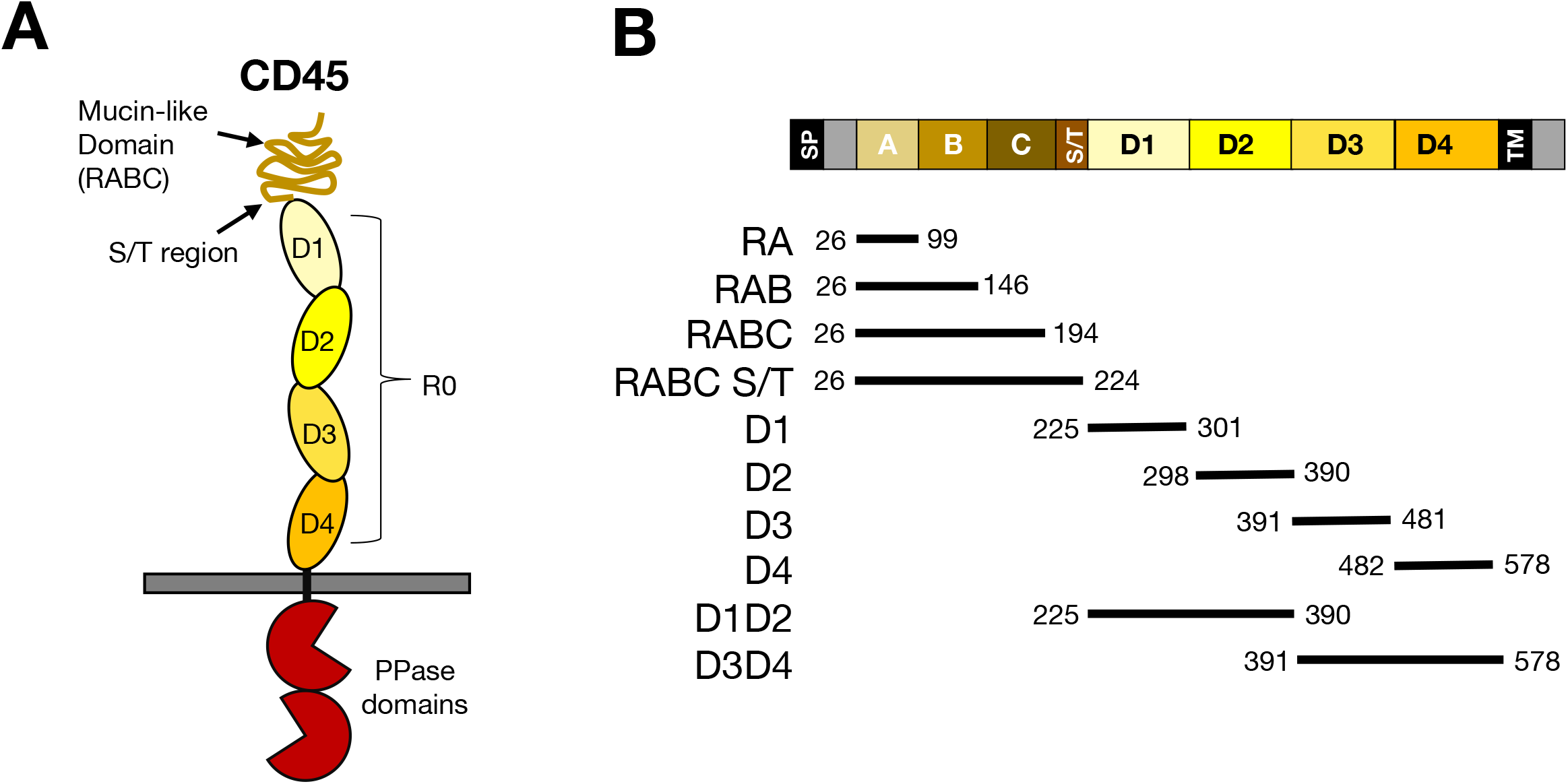
Schematic of human CD45. A) cartoon representation of human CD45, including the extracellular mucin and fibronectin like domains and intracellular phosphatase domains. B) Description of CD45 domains used for yeast display domain mapping by fusion to Aga2 protein.

In order to better elucidate which domains of human CD45 targeted by these four antibodies, we developed a yeast display system for domain mapping (Cochran et al., 2004). As shown in Fig 1B, we generated ten yeast strains containing CD45 domains fused to the C-terminus of the Aga2 protein for yeast surface expression (Fig. 1B). These constructs included each of the mucin domain exons (RA=exon 1, RAB=exon 1 and 2, RABC=exon 1,2, and 3), RABC plus S/T (RABC plus a 30 amino acid Ser/Thr rich linker between RABC and D1), each of the individual FN3-like domains (D1, D2, D3, D4), and two-domain modules (D1D2 and D3D4). The Aga2-CD45 fusions also included an internal HA epitope tag between Aga2 and CD45 as well as a C-terminal Myc epitope tag. We confirmed surface expression for each of the Aga2-CD45 fusions by HA or Myc staining (data not shown). We then used the recombinant anti-CD45 IgG molecules to stain each of the yeast strains, followed by flow cytometry to identify which strain(s) the antibodies bound to (Fig. 2A). For the VHE antibody, we found it bound to the 30 residue Ser/Thr rich linker (amino acids 194-224) between RABC and the D1 domain. The gap8.3 antibody only bound to the yeast expressing the D1D2 domain construct, suggesting an epitope that is composed of surfaces from both the D1 and D2 domains. For 4B2 and 9.4, we found both of these antibodies bound the D1 domain alone. Therefore, all four antibodies we tested bound to CD45 RO with epitopes focused in the N-terminal half of the ECD. These results are summarized in Table 1.

**Figure 2:**
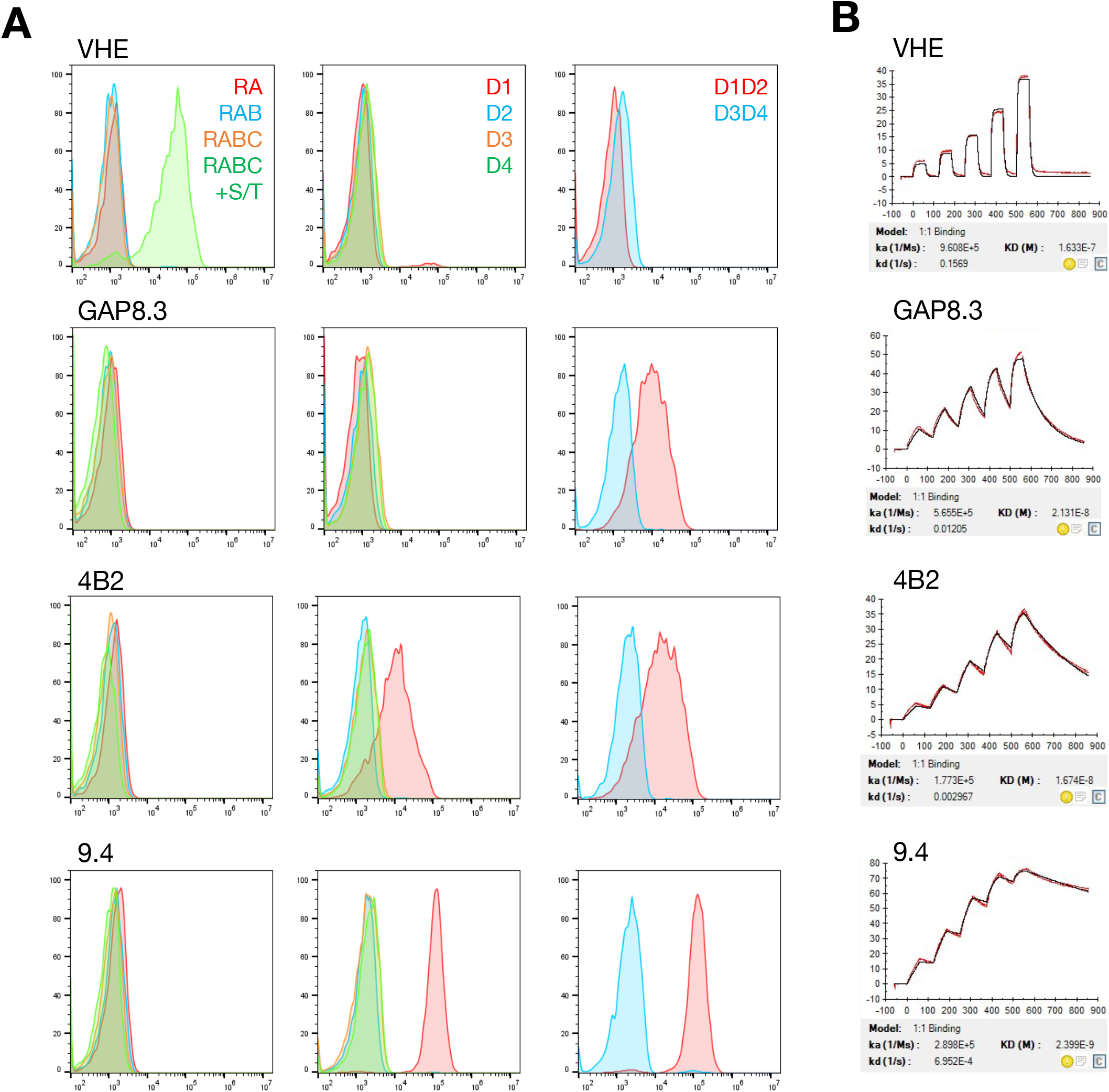
Domain mapping and affinity measurement of anti-human CD45 antibodies. A) Yeast display domain mapping of four anti-CD45 antibodies. Yeast expressing each of the domains (color coded) were stained with respective anti-CD45 IgG molecules and analyzed by flow cytometry. B) SPR analysis of CD45 binding to anti-CD45 antibodies. Human CD45 RO protein was flowed over each antibody bound to an protein A chip, ands single cycle kinetic analysis was used to determine affinity constants.

**Table 1:** Characterization of anti-human CD45 Abs

| **Clone** | **binding domain** | **IgG Affinity (K<sub>D</sub>)** | **scFv Affinity (K<sub>D</sub>)** |
| --- | --- | --- | --- |
| **VHE** | S/T | 163 nM | 260 nM |
| **4B2** | D1 | 17 nM | 12 nM |
| **9.4** | D1 | 2.4 nM | 18 nM |
| **Gap8.3** | D1D2 | 21 nM | 40 nM |

In order to identify molecules that bound to additional epitopes on human CD45, we initiated a llama VHH discovery effort using the full-length human CD45 ECD (amino acids 26-578) as an antigen. After the immunization protocol was complete, PBMCs were isolated and a phage display library constructed from the post-immune llama VHH repertoire. After one round of selection, we found excellent reactivity of the VHH library to the CD45 ECD, with >96% of the clones showing positive results in an ELISA against the CD45 immunogen. Of the 93 ELISA positive clones sequenced, we obtained 51 unique, high quality VHH sequences. Sequence analysis identified ten unique clonotypes (Fig. 3a) with varied CDR3 lengths from 10 to 27 amino acids (Fig. 3b). A subset of molecules representing each clonotype was recombinantly expressed with a monomeric Fc tag and tested for CD45RO (amino acids 194-578) binding by surface plasmon resonance (SPR). Of the ten clonotypes identified, seven bound to CD45 RO with an affinity range (K_D_) of 630 pM to 522 nM (Table 2). We then utilized our yeast display system to domain map the VHH binding sites. For labeling we used His-tagged VHH proteins incubated with the yeast and detected them with an anti-VHH antibody. We found that the seven VHH clonotypes bound across all four domains of CD45RO, with clonotypes 3, 4, and 7 targeting the D1 and D2 domains, clonotype 10 binding the D3 domain, clonotypes 2, 6, and 8 interacting with the D4 domain (Table 2). Therefore we have identified a set of VHH molecules that bind all four FN3-like domains within CD45 RO ECD.

**Figure 3:**
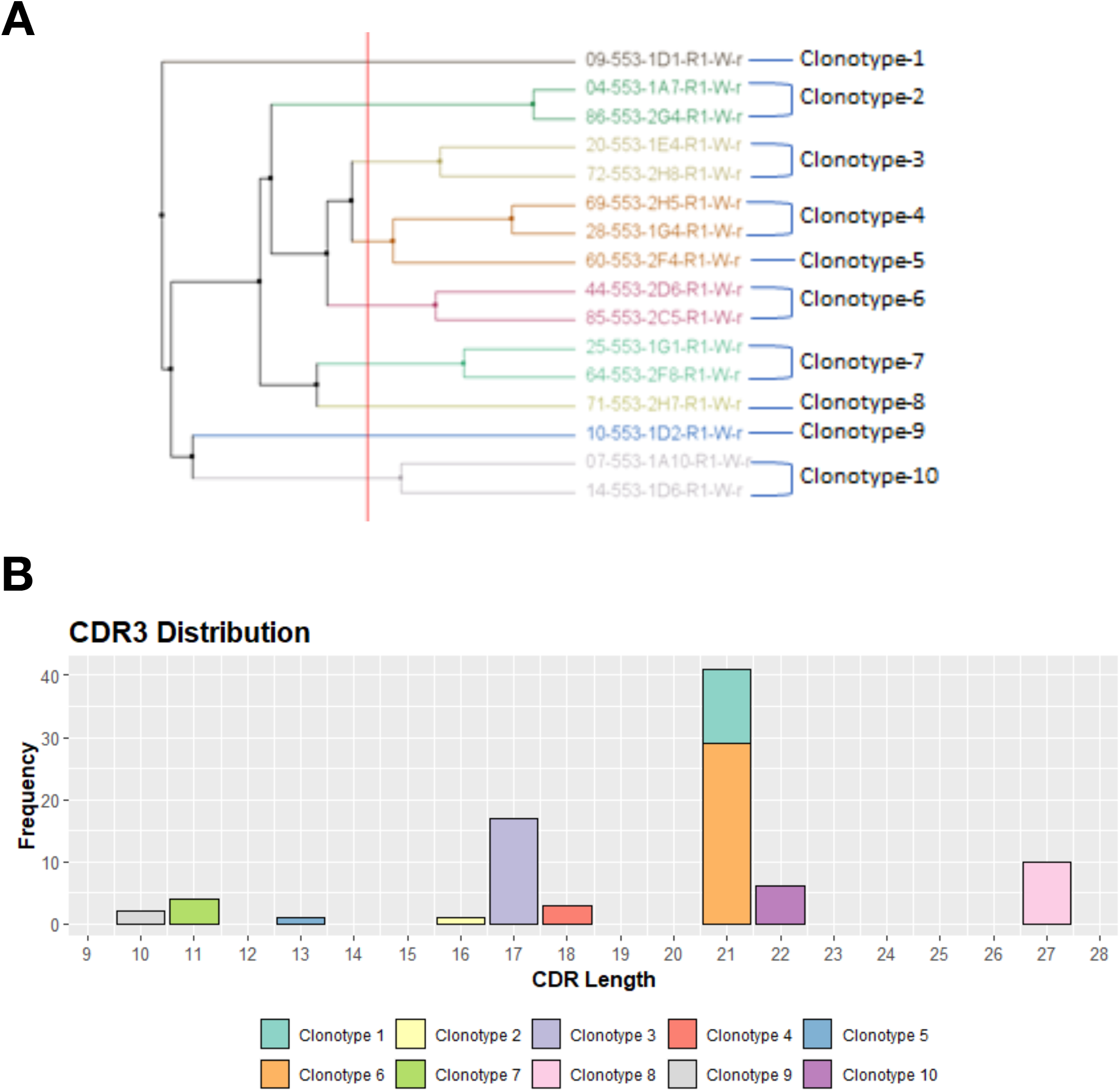
Ten anti-human CD45 VHH clonotypes show significant sequence diversity. A) Dendrogram showing the sequence relationships between VHH clonotypes identified by phage display. B) Graph showing the relative CDR3 lengths of each clonotype.

**Table 2:** Characterization of anti-human CD45 VHH

| **Clone** | **Clonotype** | **binding domain** | **Affinity (K<sub>D</sub>)** |
| --- | --- | --- | --- |
| **2G4** | 2 | D4 | 8.3 nM |
| **1E4** | 3 | D1D2 | 299 nM |
| **1G4** | 4 | D2 | 0.87 nM |
| **2H5** | 4 | D2 | 0.63 nM |
| **2C5** | 6 | D4 | 1.4 nM |
| **1G1** | 7 | D1 | 28 nM |
| **2F8** | 7 | D1D2 | 69 nM |
| **2H7** | 8 | D4 | 0.88 nM |
| **1A10** | 10 | D3 | 32 nM |
| **1D6** | 10 | D3 | 522 nM |

Our work here has shown that anti-CD45 RO antibodies available in the public domain primarily target the N-terminal half (i.e. Ser/Thr region or D1/D2 domains) of the CD45 RO ECD. This is likely due to their primary usage as tools for immune cell phenotyping by flow cytometry, which likely has selected for IgG-accessible N-terminal epitopes. In order to discover new anti-CD45-binding VHH with a more diverse epitope range, we performed a llama immunization and selection from a post-immune phage library against CD45 ECD *in vitro* to find epitopes across the entire ECD. This method has provided a VHH set with a wide range of binding affinities as well as at least five different epitopes across the CD45 RO ECD. These new VHH-based antibodies will be important tools for the study of CD45 function *in vitro* and *in vivo*.

## Methods

### Discovery of anti-human CD45 VHH

For llama immunization, the full length ECD of human CD45 (amino acids 26-578) with a C-terminal Avi-His8 tag was expressed and purified from HiFive insect cell supernatant via Ni-agarose and size exclusion chromatography in Hepes-buffered saline. A single llama was immunized according standard methods and a phage display library constructed from post-immune PBMCs. A single round of phage panning was performed against BirA-biotinylated form of the immunogen. 93 ELISA positive clones were sequenced, which provided 51 unique, high quality VHH sequences. These clones were binned into 10 clonotypes, with ultimately 7 clonotypes showing binding to the D1-D4 region of human CD45.

### Hybridoma sequencing

The anti-CD45 RO hybridomas (gap8.3, 4B2, and 9.4) were expanded using standard hybridoma culture protocols and RNA was isolated from 1×10^6^ cells. cDNA was synthesized using oligo dT primers, and 5’ RACE PCR performed using 3’ primers specific for mouse CH1. Amplicons were sequenced by Sanger sequencing.

### Yeast display epitope mapping

CD45 domains described in Figure 1 were fused to the C-term of Aga2 with an N-term HA and C-term Myc tag. Yeast constructs were expressed on the surface of yeast (*S. cervisiae* ATCC strain YVH10). Yeast were grown overnight at 30°C in synthetic selective media SD-SCAA, then induced again overnight at 20°C in SG-SCAA. Yeast were incubated with individual a-CD45 antibodies or a-CD45 VHH for 60 minutes on ice, washed, and incubated with anti-human Fc or anti-VHH Alexa647 secondary antibody as well as an anti-HA or anti-Myc Alexa488 secondary antibody. Labeled yeast were analyzed by flow cytometry on a Beckman Coulter Cytoflex and analyzed using the FlowJo software package.

### Affinity measurements

Affinities were determined using a Biacore T200 instrument by measuring K_on_/K_off_ in single cycle kinetics mode. Hybridoma clones were either reformatted and expressed in the form of the parent IgG (gap8.3, 4B2, 9.4 as murine IgG2A, VHE as human IgG1), and scFvs were expressed as human IgG1 Fc fusions. For the anti-CD45 VHH, proteins were expressed as monomeric Fc fusions. The antibody ligands were captured on a Protein A chip (GE Life Sciences) and human CD45 D1-D4 (amino acids 194-578) was flowed over the chip as an analyte diluted in HBS-EP+ buffer. Human CD45 D1-D4 was generated in house with a C-terminal 8XHis tag and expressed and purified from expi293 cells by standard Ni- and SEC chromatography methods. Kinetic affinity constants were calculated using the Biacore T200 software package.

## Notes

### Competing Interest Statement

Authors are employees of Synthekine, Inc. The authors of this paper are listed as inventors on a provisional patent application filed pertaining to the data in this paper.

